# The glycocalyx affects the mechanotransductive perception of the topographical microenvironment

**DOI:** 10.1101/2021.03.02.433591

**Authors:** Matteo Chighizola, Tania Dini, Stefania Marcotti, Mirko D’Urso, Claudio Piazzoni, Francesca Borghi, Anita Previdi, Laura Ceriani, Claudia Folliero, Brian Stramer, Cristina Lenardi, Paolo Milani, Alessandro Podestà, Carsten Schulte

**Author notes:** **<u>Co-last and corresponding authors:</u>** Carsten Schulte, Alessandro Podestà. first author (Matteo Chighizola).

## Abstract

The cell/microenvironment interface is the starting point of integrin-mediated mechanotransduction, but many details of mechanotransductive signal integration remain elusive due to the complexity of the involved (extra)cellular structures, such as the glycocalyx.

We used nano-bio-interfaces reproducing the complex nanotopographical features of the extracellular matrix to analyse the glycocalyx impact on PC12 cell mechanosensing at the nanoscale (*e.g*., by force spectroscopy with functionalised probes). Our data demonstrates that the glycocalyx configuration affects spatio-temporal nanotopography-sensitive mechanotransductive events at the cell/microenvironment interface. Opposing effects of glycocalyx removal were observed, when comparing flat and specific nanotopographical conditions. The excessive retrograde actin flow speed and force loading are strongly reduced on certain nanotopographies upon removal of the native glycocalyx, while on the flat substrate we observe the opposite trend.

Our results highlight the importance of the glycocalyx configuration in a molecular clutch force loading-dependent cellular mechanism for mechanosensing of microenvironmental nanotopographical features.

## 1. INTRODUCTION

In recent years, there has been accumulating evidence of the pivotal importance of integrin-mediated mechanosensing and mechanotransduction, *i*.*e*. the conversion of biophysical cues from the microenvironment into cellular responses. The events at the interface between the cell and the extracellular matrix (ECM) are at the base of mechanotransductive processes. At this interface, the microenvironmental biophysical cues are translated into intracellular signals, with integrin adhesion complexes (IAC) as the major relay stations. Linkages between the ECM and the actin filaments (f-actin) via integrin and talin, *i*.*e*. the formation of so-called molecular clutches, enable force transmission from the retrograde actin flow, which, in turn, is generated by actin polymerisation and the myosin-driven contraction of f-actin. These interfacial actions depend on the structural and biophysical ECM properties and regulate in an intricate manner the force loading within the molecular clutches, the maturation of IAC into signalling platforms, and cytoskeletal remodelling, which together convert into cellular responses^1,2,3,4,5,6^. The involved (extra)cellular structures in this cell/microenvironment interface, such as the ECM, the glycocalyx, the cell membrane, the IAC and the cytoskeleton, are highly complex and their sophisticated crosstalk is only partially understood.

The ECM nanotopography represents an important biophysical cue for mechanotransductive processes. The configuration of the topographical nano-three-dimensional (3D) features of the ECM sets the spatial parameters relevant for mechanosensing, such as adhesion site areas and spacing, and is therefore capable of modulating IAC formation and signalling^6,7,8,9,10,11,12,13^. Intrinsic cellular parameters can furthermore influence the cell interface with the nanotopography. The cellular cortical stiffness affects e.g. the extent to which the cell membrane can deform and contact the intervening nanostructured surface which, in turn, defines the effective accessible cell/substrate contact area at the nanoscale^14,15^. Our previous work demonstrated that the availability of activated integrins also regulates nanotopography mechanosensing and force loading^16^.

An underappreciated player in mechanotransduction is the glycocalyx, a pericellular sugar coat that is attached to the cell surface and involved in cellular adhesive processes. It is known that the glycocalyx can promote integrin clustering and influence membrane bending^2,17,18,19,20,21,22,23,24,25^. However, the studies in this context have been done on flat substrates and therefore many details about the impact of the glycocalyx on the nanotopography-dependent processes at the cell-microenvironment interface, such as force loading in molecular clutches, remain elusive; which is one of the main rationales behind this work. To this end, we studied how the glycocalyx affects the cellular capability to perceive and react to the nanotopography and impacts molecular clutch engagement and force loading (**Fig. 1**).

**Fig. 1).**
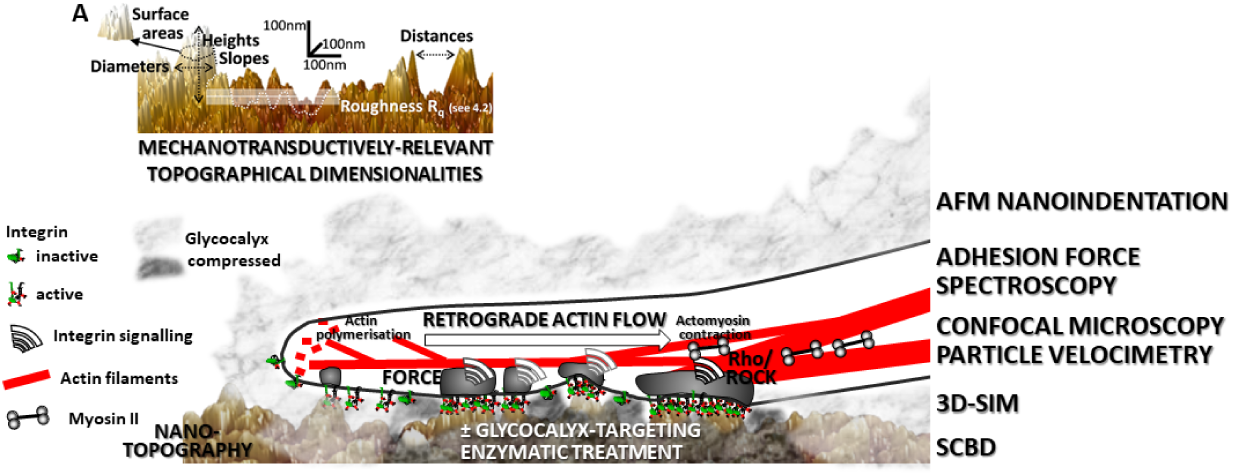
Applied approaches to address the impact of the glycocalyx on force loading-dependent mechanotransductive topography sensing at the nanoscale. The figure visualises the different mechanotransductive parameters (e.g., force loading and retrograde actin flow) and events at the cell/microenvironment interface that were scrutinised by AFM- and optical imaging-based techniques (adhesion force spectroscopy, nanoindentation, particle velocimetry, 3D-Structural Illumination Microscopy (3D-SIM)). In particular, we studied the involvement of the glycocalyx in nanotopography-sensitive and molecular clutch force loading-dependent mechanotransductive processes (by enzymatic digestion of the glycocalyx). The nanotopographical surfaces (on colloidal probes suitable for adhesion force spectroscopy, or as cell substrates as displayed here) were produced via zirconia cluster-assembling by means of supersonic cluster beam deposition (SCBD), to mimic nanostructural features cells can encounter in the ECM. **(A)** The AFM image represents the typical cluster-assembled morphology of our cluster-assembled thin films, highlighting in detail mechanotransduction-relevant topographical dimensionalities at the nanoscale.

Our approach consists in measuring the integrin-related interaction forces between living cells and suitable micro-probes or substrates, produced by gas-phase assembling of biocompatible zirconia nanoparticle^26,27^. These nano-bio interfaces are characterised by a disordered surface topography and a random distribution of asperities and valleys, whose dimensions and relative distances resemble the ECM organisation at the micro/nanoscale^16,28,29^ (**Fig. 1A**). The potential of these topographies to modulate mechanotransductive processes^9,10,13,27^, as well as cellular functioning, programme and differentiation in various cell biological contexts^9,13,30,31,32,33^, has already been documented.

Here, we aimed to dissect the role of the glycocalyx in spatio-temporal events that occur at the early stages of cell/microenvironment interaction (during IAC formation) and during nanotopography mechanosensing at long-term interaction in lamellipodia. We modelled the contact interaction geometry of cells with different nanotopographical features, as a function of cell membrane compliance with the surface, to obtain insight in the cues perceived by the cell. To analyse the early dynamics of mechanotransductive events by AFM-based adhesion force spectroscopy approaches, we challenged PC12 cells with different nanotopographical probes and manipulated the glycocalyx and the ROCK-dependent actomyosin contractility. The neuronal-like PC12 cell model was selected due to our previous studies that demonstrated how nanotopographical features robustly modulate the mechanotransductive structures and signalling in these cells^9,10,16^. We furthermore studied, by means of optical imaging techniques, the impact of the nanotopography and glycocalyx on molecular clutch engagement to the retrograde actin flow in lamellipodia.

## 2. RESULTS AND DISCUSSION

### 2.1 Structural assessment of the native PC12 cell glycocalyx

We first investigated the structural configuration of the native glycocalyx of PC12 cells, by AFM- and 3D-SIM-based techniques (**Fig. 2**) and assessed the efficiency of a comprehensive glycocalyx-targeting enzymatic treatment, consisting of a cocktail of hyaluronidase II, chondroitinase ABC, heparinase III, and neuraminidase, in removing the glycocalyx.

**Fig. 2).**
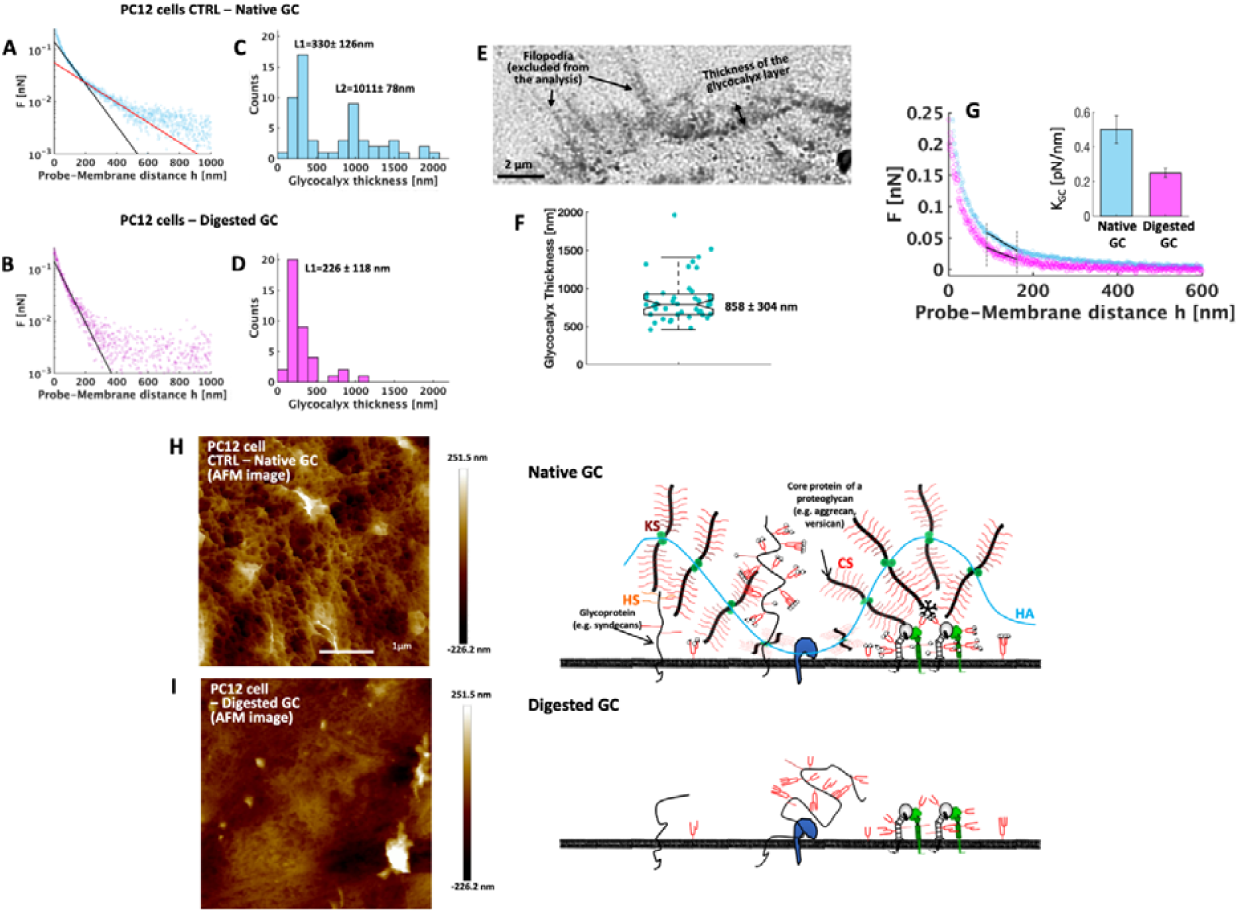
Analyses of the glycocalyx configuration by AFM-based techniques and 3D-SIM imaging before and after treatment with a glycocalyx-targeting enzymatic cocktail. **(A**,**B)** Examples of recorded Force Curves (FCs) for the measurement of the glycocalyx thickness (following the Sokolov et al. protocol^34,35,36,37^, further details can be found in SI1) on control PC12 cells with their native glycocalyx, and after enzymatic treatment for the digestion of the glycocalyx (hyaluronidase II, chondroitinase ABC, heparinase III, and neuraminidase). The force is plotted as a function of the tip/membrane distance (in semi-log scale); dots represent the experimental data and solid lines the fit using the brush model, where a linear regime highlights the entropic elasticity of the glycocalyx (GC). **(C**,**D)** Distribution of the glycocalyx thickness obtained from the experiments (one count corresponds to one cell). **(E)** Representative close-up 3D-SIM image of the periphery of a PC12 cell (on PLL), indicating the glycocalyx brush marked with wheat germ agglutinin (WGA, pixel intensities were inverted for clarity), from a plane just above the substrate used for the quantification of the thickness of the glycocalyx layer. Typical whole cell images can be found in SI – Fig. S2. **(F)** The boxplot below shows the mean and 95% confidence intervals of the glycocalyx thickness analysis from 43 such close-up zones taken from 15 different images. **(G)** Examples of two FCs with highlighted selected regions for linear fit, the inset shows the results of the effective glycocalyx stiffness. **(H**,**I)** On the left, AFM image of native PC12 cells covered with the glycocalyx (scan of the cell membrane obtained by peak force tapping technique on fixed samples) is shown (upper image), compared to the situation after enzymatic glycocalyx digestion (bottom image). On the right, the according cartoons summarise and integrate the data obtained with the AFM- and 3D-SIM-based glycocalyx analyses and visualise schematically the glycocalyx configuration in its native state and after the enzymatic treatment. HS = Heparan-Sulphate, KS = Keratan-Sulphate, CS = Chondroitin-Sulphate, HA = Hyaluronic acid.

We performed AFM indentation experiments to test the mechanical and structural properties of the glycocalyx of live PC12 cells, based on the protocol developed by Sokolov *et al*.^34,35,36,37^ (**Fig. 2 A-D**). The rescaled force curves from the untreated cells show a bilinear regime in the semi-logarithmic scale (**Fig. 2A,B**). This behaviour is commonly associated with the presence of two the glycocalyx components ^35^, one longer (red line in **Fig. 2A**), and one shorter (black line in **Fig. 2A,B**). Conversely, for the cells treated with the enzymatic cocktail, the long component was mostly (90%) removed and only the short component was present (**Fig. 2B**).

The quantification of the control PC12 cells (**Fig. 2C**) demonstrated that the long and short components of the native glycocalyx have a thickness of 1011 ± 78 nm and 330 ± 126 nm, respectively (details on the Sokolov bilayer brush model and the long and short component can be found in **SI1** and **Fig. S1-2**). The enzymatic digestion of the glycocalyx almost completely removed the long component of the glycosidic brush (**Fig. 2D**). Only in 10 % of the measured cells this long component was still present after the enzymatic treatment, and it was significantly shorter (−21.4%) with a thickness of 795 ± 150 nm. The short component was still present, but also shortened by 31.5% to a thickness of 226 ± 118 nm (**Fig. 2B,D**).

3D-SIM super-resolution imaging of live PC12 cells (**Fig. 2E, SI - Fig. S3A,B**) revealed the presence of abundant intracellular vesicles (through which glycocalyx components are transported to the surface) and a clear pericellular sugar brush around the cells. We also detected migration tracks, extracellular structures/organelles left behind by many migrating cells^38,39,40^ (**SI - Fig. S3A**). The staining of the pericellular sugar coat and the migration tracks disappeared after enzymatic treatment, with only the intracellular vesicles remaining (**SI – Fig. S3B**). Quantification of the thickness of this sugar coat at the border of the lamellipodial zones of the control cells revealed that it is 858.1 ± 304.3 nm thick, in good agreement with the values obtained by the AFM-based analysis (**Fig. 2F**).

The mechanical properties of the glycocalyx have been tested following the so-called “mechanical spring model” approach (for details, see Methods **4.3.3**). The results of the analyses demonstrate a glycocalyx stiffness of K_GC_= 0.50 ± 0.08 pN/nm that decreased ∼50% after the enzyme digestion to K_GC_= 0.24 ± 0.03 pN/nm (**Fig. 2G**).

Imaging of fixed PC12 cells (with or without enzymatic treatment) by AFM showed that the native cell surface was characterised by porous, reticular and filamentous structures (**Fig. 2H**), which became smooth after the enzymatic treatment with few remaining agglomerates (**Fig. 2I**).

Altogether, these findings confirmed the presence of a substantial glycocalyx layer, approximately 1 μm thick, around native PC12 cells and the efficacy of the glycocalyx disruption by the enzymatic treatment.

### 2.2 Modelling of the surface accessibility as a function of cell membrane compliance to the nanotopographies

In this work, we used three different topographies (typical representations of the morphological features are shown in **Fig. 3A and SI3 – Fig. S4-5**): a featureless flat zirconia substrate (flat-Zr, produced by ion gun sputtering) with a roughness parameter R_q_ <1 nm, and two nanostructured zirconia substrates with a R_q_ = 15 nm (ns-Zr15) and R_q_ = 20 nm (ns-Zr20); the latter two produced by SCBD (fabrication details in **4.1-2**). The structure and morphology of the nanostructured films result from the random stacking and aggregation of impinging nanometric particles of ZrO_2_ and closely mimic nanotopographical features that cells encounter in ECM with spatial parameters that are relevant for mechanotransduction ^16,28,29^.

**Fig. 3).**
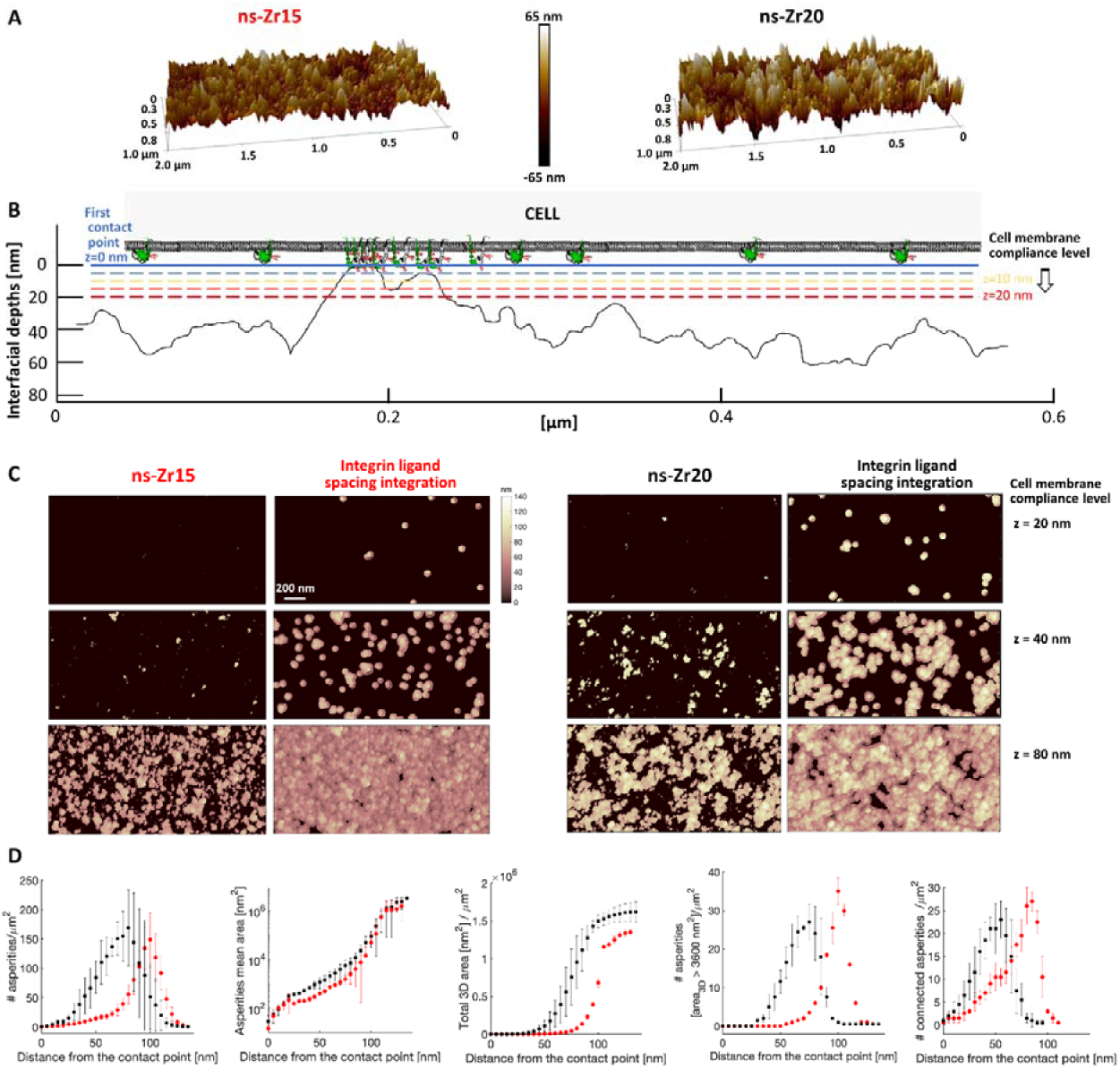
Modelling of the accessibility of nanotopographical in dependency of cell membrane compliance to the morphology. **(A)** These images demonstrate representations of the morphological features (in 3-dimensional views) of the different topographies (ns-Zr15 and ns-Zr20, produced by SCBD) that have been tested in the experiments (an example of the featureless flat-Zr, produced by Ion Gun sputtering, can be found in **Fig. SI-S4**). **(B)** Representative profile of a cluster-assembled zirconia substrate, with the cell membrane (and embedded integrins) above and the first thresholds for membrane compliance used for the analysis. **(C)** The images illustrate the accessible nanotopographical features at different cell membrane compliancy levels (20, 40, 80 nm of interfacial depth) for ns-Zr15 and ns-Zr20 (respective left images). The respective right images show the situation, if asperities that are in 60 nm adjacency (ligand spacing threshold) are converged. **(D)** Statistics of the determined parameters that are relevant for cell adhesion and mechanotransduction: # Asperities, Asperities mean area, Total 3D area, # of asperities with > 3600 nm^2^ (minimal adhesion unit), # of connected asperities. Red symbols = ns-Zr15, black symbols = ns-Zr20.

Compared to flat substrates, the three-dimensionality of the nanotopographical surfaces adds a critical level of morphological complexity to cell adhesion processes (**Fig. 1A**). The effective cellular contact area with a given nanotopography can vary decisively as corollary of the capability of the cell membrane and its embedded adhesion receptors to access, or not, the bottom parts of nanotopographical asperities. This is still a rather underestimated aspect, even though it was recently found by Park *et al*.^14,15^ that different compliance of the cell membrane with a nanotopography can impact important (patho)physiological cellular processes, such as cell migration and behaviour of metastatic cells. Our previous characterisations demonstrated that the dimensionalities of the nanotopographical asperities produced by SCBD^9^ influence mechanotransductive processes^9^.

Moreover, work by Paszek *et al*.^18,41^ has shown that the glycocalyx, as structure in the cell/microenvironment interface, forms a steric barrier for integrin/ligand binding outside of already established adhesion sites with the substrate, due to its bulkiness and compression^1,2^, and can impact the membrane bending^23^. However, these studies were performed in the context of flat substrates. It can be hypothesised that, also in the case of nanotopographical substrates, the presence of a long glycocalyx (*e*.*g*., hundreds of nm for the PC12 cells, **Fig. 2C,E,F,H**) will impede the access of adhesion receptors to lower areas of asperities. Our measurement of the glycocalyx stiffness (**Fig. 2G**), supports this assumption. Indeed, in presence of their native glycocalyx, PC12 cells interact predominantly with the apical part of the nanotopographical asperities^9^.

However, very little is known on how these two essential factors at the cell/microenvironment interface, the nanotopography and the glycocalyx, affect nanoscale mechanotransductive processes in combination.

We were interested in modelling the accessibility of nanotopographical asperities for the cell in dependency of cell membrane compliance with the substrate, and how this would affect the corresponding configuration of the cell/substrate interface (**Fig. 3B**). We analysed morphological AFM maps of ns-Zr15 and ns-Zr20 by measuring, at different interfacial depths, several parameters that are significant for cell adhesion and mechanotransduction (**Fig. 3C,D**). Into this model, we incorporated known mechanotransduction-relevant factors, such as ligand spacing and minimal adhesion unit (for details on the model, see **Methods 4.3.5**). For ligand spacing, we set a value of 60 nm, based on the identified threshold that regulates the formation of IAC^42,43,44^. In addition, Changede *et al*.^45^ found that nanocluster bridges of unligated integrins can form between adjacent (tens of nm) nanometric adhesion sites which alone are not sufficient to sustain IAC maturation. With respect to the minimal adhesion unit, we determined the number of those asperities at the different interfacial depths that are characterised by a 3D surface area of at least 3600 nm^2^, because it has been shown that, at least, four integrin binding sites within 60 nm can serve as such a minimal adhesion unit that promotes IAC maturation^45,46,47,48^.

The objective behind this modelling was to get an idea of the kind of nanotopographical cues the cell might perceive at different levels of compliancy (**Fig. 3C**). The morphological differences between nanotopographical substrates have the potential to shift spatial mechanotransduction-relevant factors, as we have previously shown in a simpler model^9^. However, the disordered nature and complex geometrical properties of these nanostructured asperities prompt an in-depth analysis considering also the cell membrane compliancy, because integrin clustering depends on the local arrangement of adhesion sites at the nanoscale rather than on the global average ligand density^49^.

The number of accessible asperities initially increases for both conditions, ns-Zr15 and ns-Zr20, when reaching deeper interfacial depths, but deeper compliancy levels are necessary for ns-Zr15. Also, the mean area of these asperities and the total 3D surface area is lower for ns-Zr15 than for ns-Zr20 at most compliancy levels (except for very low compliancy) (**Fig. 3C,D**). Consistently, after the integration of the ligand spacing factor, the convergence of asperities which are in a 60 nm adjacency can only be achieved at much higher compliance level for many asperities in the ns-Zr15 condition, as well as the spatial requisites for minimal adhesion unit (**Fig. 2B, Fig. 3C,D**).

These results demonstrate, as expectable, that stronger cell membrane compliance to the nanotopographical surface can provide an increased number of interaction sites with spatial conditions that are suitable for integrin clustering and IAC maturation. These specific effects are, however, highly dependent on the given roughness of the nanostructured zirconia substrate. Indeed, a counterintuitive property of these cluster-assembled zirconia interfaces is that the rougher ns-Zr20 substrates display properties, in terms of mechanotransduction-relevant parameters, that are closer to the flat surface situation (which has no spatial restrictions) over a wide range of compliance levels; *i*.*e*., a potentially less strict confinement of interaction areas for integrin adhesion. The model predicts instead a stark difference of mechanotransduction-relevant parameters between the low (corresponding to the presence of native glycocalyx) and higher compliancy level (without glycocalyx) for the ns-Zr15 substrate.

The model suggests that the nanotopographical cues appear as a kind of “3D QR code” (**Fig. 3C**) whose read-out will be affected by the ability of the cell membrane to reach lower parts of the asperities. Divergent levels of cell membrane compliance will provide different spatial surface information to the cell, with the potential to influence mechanotransductive nanoscale processes, such as force loading and molecular clutch engagement.

### 2.3 Adhesion Force Spectroscopy for the characterisation of the impact of the nanotopography and glycocalyx on early integrin-related interactions

It has been demonstrated previously that spatial restrictions of the cellular adhesion site dimensions at the nanoscale (as in the case of ns-Zr15) keep IAC at the focal complex size and diminish stress fibre development, compared to featureless flat substrates on which the IAC mature to FA and stress fibres form^9^. Recently, we have also shown that interaction with these specific nanotopographical features can cause excessive force loading in integrin-mediated nanometric adhesion sites during nascent adhesion formation, leading to their disassembly, regulated by availability of activated integrins^16^.

Here, we analysed how the adhesion and force loading dynamics towards different nanotopographical features change in the presence or absence of the glycocalyx (**Fig. 4**).

**Fig. 4).**
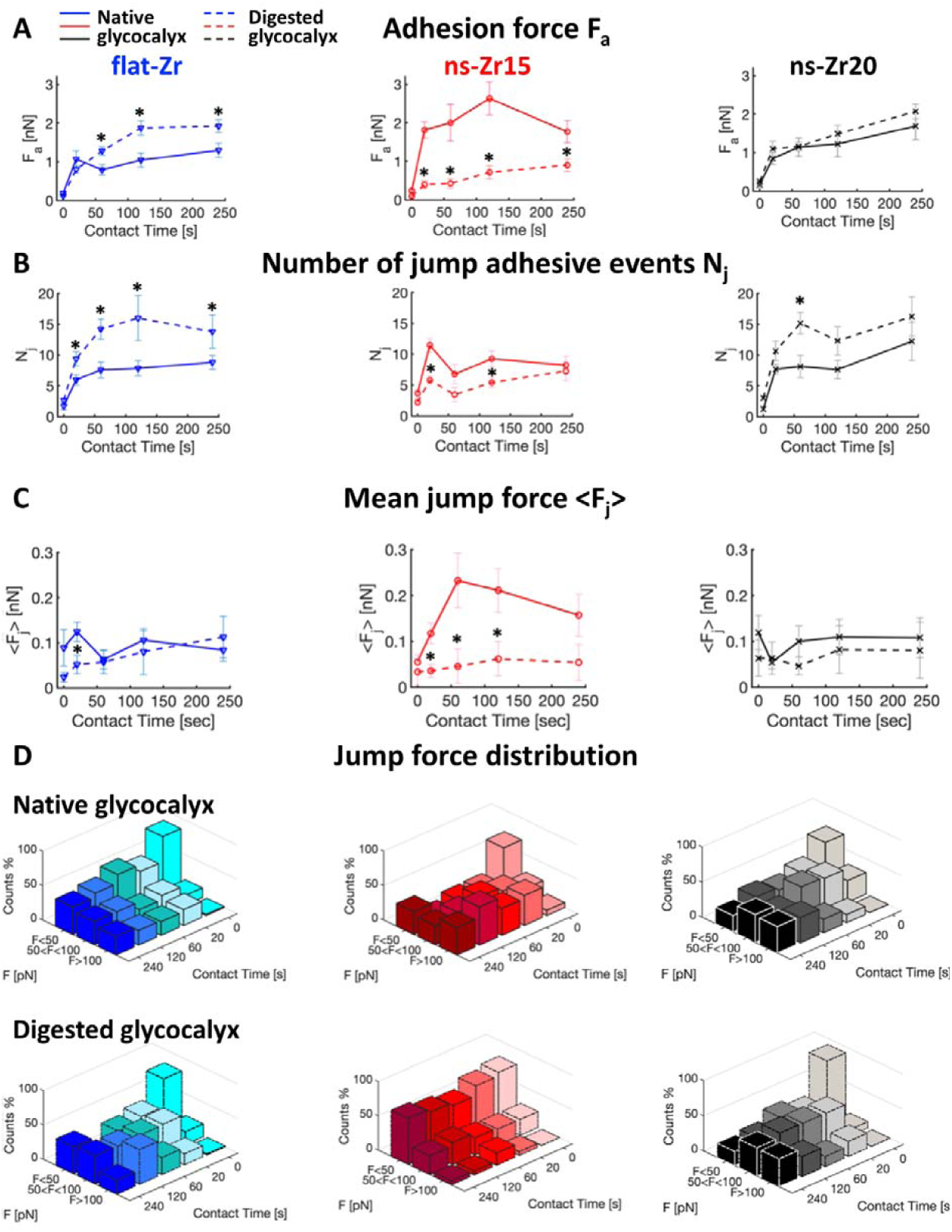
Differences in the nanotopography can induce specific adhesion dynamics that strongly depend on the glycocalyx configuration. The panel shows the results of the adhesion force spectroscopy measurements for probes with flat-zirconia films devoid of nanotopographical features (flat-Zr, blue lines or bars), and with nanostructured cluster-assembled zirconia films with a roughness R_q_ of 15 nm (ns-Zr15, red lines or bars), or 20 nm (ns-Zr20, black lines or black/grey bars), in the presence of the cell’s native glycocalyx (solid lines or bars), or after glycocalyx removal by enzymatic digestion (dashed lines or bars with border lines). The measurements were taken at 5 different cell-probe contact times (0 s, 20 s, 60 s, 120 s, and 240 s). The parameters presented in this graph are **(A) Maximum adhesion force F**_**a**_, **(B) Number of jump bonds N**_**j**_, **and (C) Mean strength of jump bonds <F**_**j**_**>** (Work W and Number of tether bonds N_t_ can be found in **SI4 – SI Fig. S6**). The error bars represent the effective standard deviation of the mean details in **4.3.4**). Asterisks indicate significant differences between control and glycocalyx-targeting enzymatic treatments. **(D)** The bars show the temporal evolution of the jump force distributions for the different experimental conditions, allocating the forces into 3 categories, i.e. low forces <50 pN, intermediate forces, 50-100 pN, high forces >100 pN (the original distributions of bond strength can be found in the **SI4 – SI Fig. S7**, to see the dispersion of higher forces.

We therefore performed adhesion force spectroscopy^16^ on PC12 cells with their native glycocalyx and after treatment with glycocalyx-targeting enzymes. We used colloidal probes coated with three different topographies: flat-Zr, ns-Zr15, and ns-Zr20 (typical representations of the morphological probe features are shown in **SI3 - Fig. S5**). We selected five cell-probe contact times covering different stages of the critical window of nascent adhesion formation towards maturation to focal adhesions^50,51^, *i*.*e*. 0 s, 20 s, 60 s, 120 s, and 240 s.

From these measurements we derived information on the maximum adhesion force F_a_ (**Fig. 5A**), the number of jumps N_j_ (**Fig. 5B**), the mean jump force <F_j_> (**Fig. 5C**), the distribution of jump force (**Fig. 5D**), the work W (**SI4 – SI Fig. S6A**), and the number of tethers N_t_ (**SI4 – SI Fig. S6B**). Jump events are usually associated with membrane adhesion receptors that are anchored to the cytoskeleton, as for integrins engaged in molecular clutches^52,53,54,55^, while the tether events are attributed to receptors that are not bound to the cytoskeleton^56,57^.

**Fig. 5).**
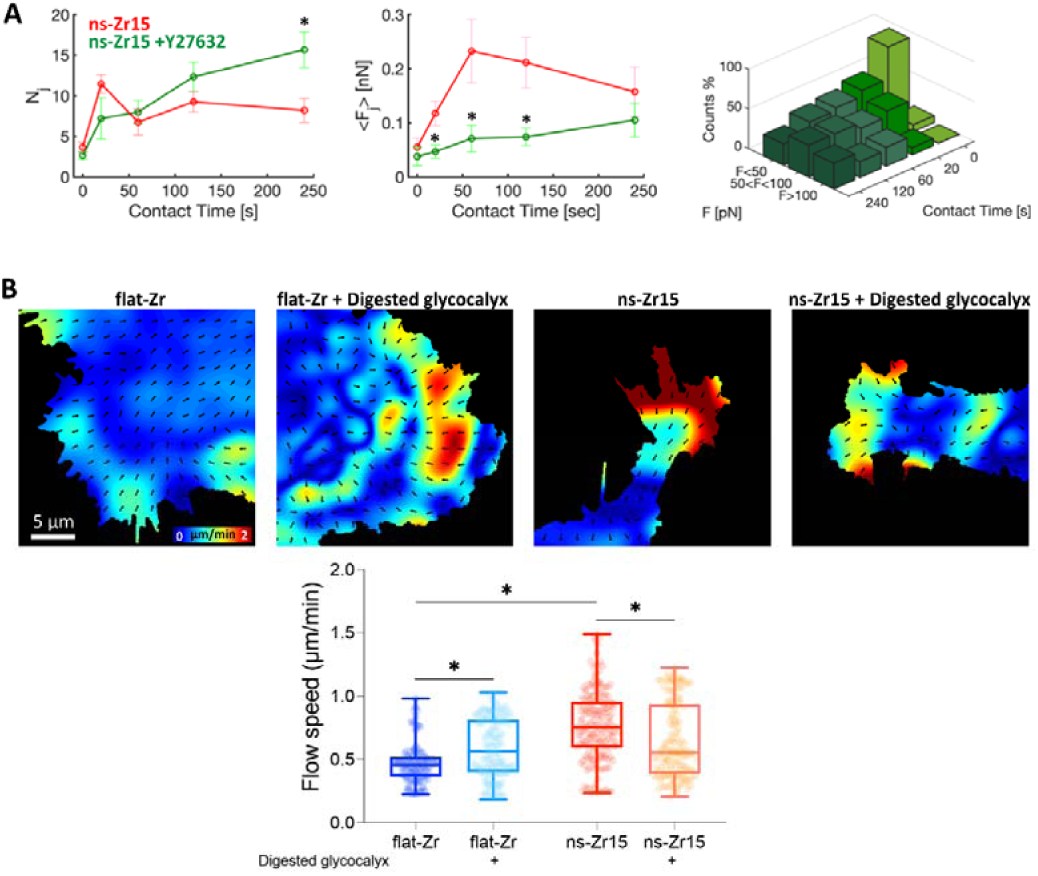
The nanotopography and the glycocalyx configuration affect the lamellipodial molecular clutch engagement and actin dynamics. **(A)** The graphs summarise the evolution of N_j_, <F_j_> and jump force distribution comparing untreated PC12 cells interacting with ns-Zr15 (red lines, reproduced from **Fig. 4B,C**), or treated the ROCK inhibitor Y27632 (10 µM, green lines or bars, for ns-Zr15 Native glycocalyx jump force distribution, compare with **Fig. 3D**). The error bars represent the effective standard deviation of the mean (details in **4.3.4**). **(B)** The panel shows representative actin flowfield images obtained after Particle Image Velocimetry (PIV) of live cell recordings of PC12 cells (transfected previously with LifeAct™-mCherry to visualise the actin dynamics) in the different experimental conditions. The confocal recordings had a frame rate of 0.5 images/sec. The graph below shows the according PIV-based quantification of the actin dynamics. The boxplots show medians, 25^th^ and 75^th^ percentile as box limits. 198-234 frames from 11-14 cells/lamelllipodial zones were quantified. A Kruskal-Wallis statistical test was applied with multiple comparisons (flat-Zr without vs. with glycocalyx digestion, ns-Zr15 without vs. with glycocalyx digestion, flat-Zr vs. ns-Zr15 without glycocalyx digestion, flat-Zr vs. ns-Zr15 with glycocalyx digestion). * p < 0.0001.

#### 2.3.1 Nanoscale differences in the topography modulate the temporal adhesion and force loading dynamics

We first analysed the impact of the three different topographies in the presence of the native glycocalyx (solid lines in **Fig. 4**). The results for flat-Zr and ns-Zr15 closely reproduced the outcome of our previous work^16^. The adhesion force F_a_ was generally low in the flat-Zr condition, and the number of jumps N_j_ increased progressively until reaching a plateau. A different situation was found for the ns-Zr15 condition, where F_a_ reached higher values and N_j_ fluctuated heavily, *i*.*e*. first increasing strongly at 20 s and then showing a drop towards 60 s. This was reflected in a significant divergence of mean jump strength <F_j_> for the two conditions. Whereas this value had only minor variations in the flat-Zr condition, a rise to higher values in the ns-Zr15 condition with a maximum at 60 s was observed. Generally, the range of N_j_ values at later time points are in good accordance with the potential adhesion sites predicted by the model after integrating the ligand spacing factor (compare **Fig. 3C,D** with **Fig. 4B**).

Interestingly, the values for ns-Zr20 condition were instead akin to the flat-Zr condition, and not to the ns-Zr15 one, in agreement with the model predictions: a smooth increase in F_a_ with low absolute values comparable to those obtained on flat-Zr, and a constantly low <F_j_>. The values for W and N_t_ wavered in a similar range for all three conditions (**SI4 – SI Fig. S6**, solid lines). A dissection of the jump force distribution (**Fig. 4D**) showed for flat-Zr that most jump forces were in the <50 pN category at all time points, with only a minor increase of intermediate (50-100 pN) and higher forces (>100 pN) over time. For ns-Zr15, there was instead an early shift (starting already at 20 s) towards the higher forces, *i*.*e*. >100 pN, even with appearance of some very high forces. The ns-Zr20 condition was intermediate, the jump force distribution was more equilibrated than in the other two conditions, with the 50-100 pN category being the predominant one for most time points (except 120 s).

These data show that small differences in nanoscale topographical features can have a significant effect on early cellular adhesion parameters and interfacial force-related mechanotransductive processes at the cell membrane level.

#### 2.3.2 The configuration of the glycocalyx impacts on the mechanosensing of the nanotopography

After digestion of the glycocalyx, some interesting and divergent effects could be observed (dashed lines in **Fig. 4** and **SI4 – Fig. S6**).

For the ns-Zr15 condition, the glycocalyx removal had a strong impact on several parameters: it reduced F_a_, N_j_, <F_j_> and W to values that were actually very similar to those we observed for the flat-Zr condition with the native glycocalyx, or even below, from 20 s onwards.

For the flat-Zr condition, the removal of the glycocalyx had instead the opposite effect for F_a_ and N_j_, leading to increased values that are comparable to the ns-Zr15 situation with the native glycocalyx or even higher, whereas W and <F_j_> (except 120 s) remained basically unchanged. For ns-Zr20, the variations due to the glycocalyx-targeting enzymatic treatment were more modest, the W (for the 120 s and 240 s time points) and N_j_ (at 60 s) increased at certain time points, whereas F_a_ and <F_j_> did not alter. The removal of the glycocalyx had a drastic effect on the jump force distribution (**Fig. 4D**) in the ns-Zr15 condition, as most of the jump forces were found in the <50 pN category for all time points. A minor fraction of forces was in the 50-100 pN category, whereas higher forces were practically absent. This was an almost complete inversion compared to the untreated cells interacting with this nanotopographical surface. The glycocalyx-targeting treatment had instead little impact on the jump force distribution in the flat-Zr condition (slight shift to more intermediate forces) and there was almost no alteration for ns-Zr20 condition.

An interesting aspect was also observed with respect to the most probable jump forces at 0 s. The values for these first pristine interactions forming between integrins and substrate were reduced after glycocalyx removal in all topographical conditions (**Table 1**, see also **SI4 – SI Fig. S7**). This is congruent with the mechanical loading of integrins due to the adjacent compressed glycocalyx shown by Paszek *et al*.^18,41^, and our data on the glycocalyx stiffness (**Fig. 2G**).

**Table 1.** Most probable jump forces at 0 s. The table shows the most probable jump forces at 0 s for the different experimental conditions.

| Topography/experimental condition | Native glycocalyx | Digested glycocalyx |
| --- | --- | --- |
| flat-Zr | $44 \pm 16$ pN | $25 \pm 18$ pN |
| ns-Zr15 | $39 \pm 16$ pN | $28 \pm 10$ pN |
| ns-Zr20 | $43 \pm 17$ pN | $33 \pm 11$ pN |

These results show that the glycocalyx has a decisive influence on the early adhesion dynamics towards nanotopographical features and that this influence depends on the nanometric details of the topography, as evident especially for the case of ns-Zr15 (which has been shown to strongly modulate IAC-mediated mechanotransductive processes and signalling in PC12 cells^9,10,16^). Indeed, the impact of glycocalyx removal can strongly differ in dependency of the particular nanotopographical conditions, as also suggested by the model.

### 2.4 The glycocalyx configuration affects nanotopography-sensitive molecular clutch engagement to the retrograde actin flow

The forces that drive the force loading within molecular clutches derive from the retrograde actin flow in lamellipodia, which, in turn, is generated by actin polymerisation and actomyosin contraction, regulated by Rho/ROCK signalling^1,2,5,6,58,59,60,61,62,63,64,65,66^. An inverse relationship between actin retrograde flow velocity and the traction forces has been demonstrated during transition from nascent adhesion to FA, i.e. IAC maturation causes actin flow deceleration by molecular clutch engagement and reinforcement^60,61,62,64^.

For flat substrates, it is known that the initial integrin clustering in nascent adhesions is independent of actomyosin-driven forces^60,67^. The force loading in molecular clutches, IAC growth to FA, and stress fibre formation, instead strongly depend on actomyosin contraction.

To understand whether the observed nanotopography-sensitive force loading dynamics in the nascent adhesions depend on actomyosin-driven contractility, force spectroscopy measurements with the ns-Zr15 probes on PC12 cells treated with the ROCK inhibitor Y27632 have been performed. The results revealed a clear change of the adhesion dynamics and forces after the Y27632 treatment (**Fig. 5A**). Differently from the ns-Zr15 of untreated cells, N_j_ progressively increased over time after the inhibition of the actomyosin contraction, reaching particularly high levels at 120 s and 240 s. Compared to the untreated ns-Zr15 condition, <F_j_> was strongly reduced for the time points in the critical nascent adhesion formation window (20 – 120 s), with appearance of fewer high forces. Also, the most probable jump force at 0 s was reduced (25 ± 14 pN, compare to the value of untreated cells reported in **Table 1**, *i*.*e*. 39 ± 16 pN). These data indicate the importance of actomyosin-driven forces for the specific ns-Zr15 adhesion force dynamics.

Lamellipodia are integrators of the biophysical cues that are present in the local cellular microenvironment. To that effect, the lamellipodial retrograde actin flow velocity is a good indicator for the overall force loading-dependent molecular clutch engagement induced by a specific substrate^1,2,4,6,59^. Basically, the retrograde actin flow changes as a function of the configuration and overall maturation status of the IACs inside the lamellipodia. To test how the nanotopographical substrate cues and the glycocalyx configuration are integrated in a more long-term and comprehensive adhesion scenario, we measured the retrograde actin flow speed (and accordingly the extent of molecular clutch engagement), by recording the actin cytoskeletal dynamics (visualised by LifeAct™ transfection) of PC12 cells that interact with flat-Zr or ns-Zr15 substrates in the presence or absence of the glycocalyx.

Particle image velocimetry (**Fig. 5B and corresponding videos**) demonstrated that the lamellipodial actin flow dynamics were significantly faster in the cells plated on ns-Zr15, compared to flat-Zr. This is in good accordance with the fact that on ns-Zr15 the IAC have focal complex dimensions due to the nanotopography-imposed spatial restriction for integrin nanocluster formation, whereas there is focal adhesion formation on flat-Zr^9^. This nanotopography-dependent difference in actin flow speed equals when the cells were treated with the glycocalyx-targeting enzymes, meaning that the actin flow speed increased on flat-Zr and decreased on ns-Zr15.

Notably, the opposite effect on the retrograde actin flow to glycocalyx removal that was observed between flat-Zr and ns-Zr15, which represents a more comprehensive adhesion effect, is congruent to the phenomena seen for the nascent adhesion force dynamics in the force spectroscopic measurements. These data demonstrate that the nanotopography mechanosensing relies on Rho/ROCK signalling-regulated actomyosin contraction and that molecular clutch engagement is nanotopography-sensitive in a glycocalyx-dependent manner.

For flat-Zr, the effects of the glycocalyx removal are in good accordance with the paradigm established by Paszek *et al*.^18,41^, *i*.*e*. a bulky glycocalyx layer promotes integrin clustering by funnelling of active integrins to an initial integrin/ligand binding site (“kinetic trap”) and by applying additional tension due to upward force caused by the adjacent glycocalyx compression. The force spectroscopy results suggest that the glycocalyx’ steric repulsion indeed strengthens the initial bonds, and that, in the absence of the glycocalyx, the cells create more bonds with the microenvironment. However, the increase in the actin retrograde flow speed after glycocalyx removal indicates that abolition of the glycocalyx-dependent effects reduces the molecular clutch engagement and integrin clustering on the flat substrate.

Nanotopographical features apparently add another level of complexity, leading to particular glycocalyx-sensitive phenomena that are subject to the exact topographical configuration, as they are specific for the ns-Zr15 condition. In the presence of the native glycocalyx, the initial nascent adhesion formation, which is independent of actomyosin contraction, seems to be promoted in the ns-Zr15 condition (high N_j_ towards 20 s). However, the excessive force loading-dependent events (increase of <F_j_> and drop of N_j_, which are dependent on actomyosin contraction) lead eventually to a lower molecular clutch engagement to the retrograde actin flow (higher speed compared to the flat-Zr condition). Concordantly, we know from our previous work that the IAC are of much smaller nanometric dimensions (focal complex size) on ns-Zr15, compared to the micrometric FA on flat-Zr^9^. The deceleration of the retrograde actin flow speed, after glycocalyx removal, insinuates instead an increased molecular clutch engagement compared to control ns-Zr15 condition. Glycocalyx removal, by enabling easier membrane bending around the asperities due to the omission of the repulsive forces of the compressed glycocalyx^2,18^, has the potential to increase the accessible surface contact area^14^ (**Fig. 3C,D, Fig. 6**). The differences between ROCK inhibition and glycocalyx removal for the ns-Zr15 situation, especially regarding N_j_ (compare **Fig. 5A** and **Fig. 4B**), indicate a complex contribution of the glycocalyx configuration in nanotopography sensing.

**Fig. 6).**
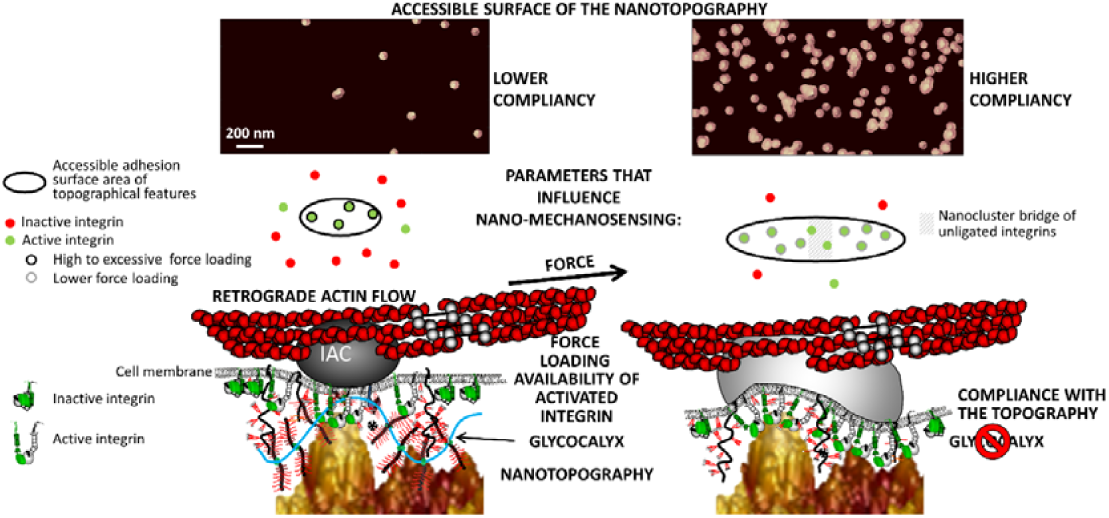
Model for the way the glycocalyx affects force loading-dependent mechanosensing of the nanotopography. The figure graphically summarises the results of this work, integrating some insights from the literature published by us and others^1,2,16,18,45^, and visualises how the glycocalyx configuration might modulate the cellular perception of nanotopographical features the cells interacts with. The presence or absence of the glycocalyx has the potential to influence the membrane bending and compliance with the nanotopography which changes the surface area of the asperities that is accessible for integrin adhesion complexes. This alters the integrin clustering and the force distribution/loading within the molecular clutches of nascent adhesions.

## 3. CONCLUSIONS

Our results shed some light on how the glycocalyx affects the cellular perception of nanotopographical features and impacts mechanotransductive events at the nanoscale.

Altogether, our data clearly demonstrate that the glycocalyx decisively modulates the dynamics of cellular adhesive and mechanotransductive processes in a nanotopography-dependent way, particularly regarding to the number and structuring of adhesion sites at the nanometric scale and the force distribution within them. We show that the nanotopography and glycocalyx represent crucial factors within the cell/microenvironment interface that must be considered in mechanotransduction in an interdependent manner. The configuration of the glycocalyx and further intrinsic cellular parameters, such as the type and availability of activated integrins, and the extent of engagement to actomyosin-driven forces, control how a cell perceives the nanotopographical features it interacts with. We have demonstrated, for PC12 cells, how manipulations of the glycocalyx, integrin activation and actomyosin contraction change the topography mechanosensing at the nanoscale. As a function of these intrinsic (extra)cellular factors, the same nanotopography can convey diverse cues to the cell (**Fig. 6**).

Di Cio *et al*.^68^ found, similar to our results, that cells became insensitive to the width of nanotopographical features (in their case nanofibers) upon ROCK inhibition. They suggest an actin architecture-dependent long-range, rather than FA-dependent, mechanism for nanotopography sensing, based on actin network collapsing into foci.

Differently to Di Cio *et al*., we here propose a molecular clutch force loading-dependent mechanism for nanotopography sensing, with a decisive involvement of the glycocalyx. The effect of the ROCK inhibition on the nanotopography sensing (through its impact on the force loading), *i*.*e*. that the cells become blind for the nanotopographical differences, has some similarities to the role of actomyosin contraction (by sarcomere-like contractile units) and force production in the rigidity sensing mechanism described by Wolfenson *et al*.^69,70^. They have shown that the rigidity sensing occurs by actomyosin-driven stepwise contractions. These contractions in themselves are independent of the substrate rigidity and non-mechanosensitive^71^, but a force threshold of around 20-25 pN has to be reached to enable IAC reinforcement^69^.

Albeit the substrates we used here are rigid, we have shown that these nanotopographies can directly affect the extent of integrin clustering and the IAC size at the nanoscale^9^, as well as the force loading in molecular clutches^16^. On ns-Zr15, the effects on mechanotransduction resemble what can be observed on soft substrates^1,2,58,72^. Considering the work of Oria *et al*.^59^ that points out the importance of the force loading in molecular clutches for spatial ligand sensing, the rigidity and nanotopography mechanosensing are likely to be interdependent. Our data strongly indicate that the glycocalyx and the nanotopography configuration must be taken into account, as alterations therein can change the whole setting in the cell/microenvironment interface.

The dynamic biophysical cue-dependent events in the cell/microenvironment interface exhibit an intriguing complexity. Understanding the mentioned interdependencies in more detail will be a challenging and very interesting field for future investigations and it is of high biomedical relevance, in particular with respect to cancer cell biology^15,21,24,70,73,74,75^.

It can be speculated that these general mechanisms in the cell/microenvironment interface might be similar for many cell types. Specific adhesion and force loading dynamics towards nanotopographies will certainly differ, depending on the cell type-specific glycocalyx thickness and composition, integrin expression levels, integrin trafficking, the contractile machinery, cytoskeletal organisation and mechanics. One of our future objectives will be to dissect such differences (or find similarities in patterns) between cell types, and/or between healthy and pathophysiological situations.

The potential implications of these cell/microenvironment interfacial processes for pathophysiological situations become evident from the fact that aberrations in the components along the mechanotransductive sequence have been associated with various diseases^76^, particularly to cancer and metastasis, as well as neurodegenerative diseases^21,73,74,75,77,78^. Examples are abnormal rearrangements and composition of the ECM^79,80,81^ and glycocalyx^82^, as well as dysregulation in IAC-related genes^83,84^, or in the contractile machinery^75^. Recently, it has been shown that a glycocalyx-dependent potentiation of an integrin-mediated mechanosignalling feedback loop might be causally involved in the primary brain cancer glioblastoma multiple^85^.

Here we concentrated on direct integrin-mediated mechanotransduction, but these results could have also broader relevance in the context of other surface receptors (such as syndecan-4^86^, CD44^87^, uPAR^64,88^), receptor clustering in synaptic perineuronal nets^89^, and curvature-sensing proteins (such as FBP17)^90^.

Obtaining a deeper understanding of the nanoscale mechanotransductive events at the cell/microenvironment interface is thus essential. It is furthermore highly attractive from a biomedical, diagnostic and therapeutic perspective because the cellular outside and surface are easier to access than intracellular processes. This manifests in a recent re-evaluation and emergence of interest in potential ECM- and integrin-targeting drugs, or more general, so-called mechanotherapeutics^77,81,91,92^.

## 4. MATERIALS AND METHODS

### 4.1 Fabrication and calibration of the colloidal probes

The procedure for the fabrication of colloidal probes (CP) is based on the approach described elsewhere^93,94^. The most important highlights of the protocol and the advantages of the probes that can be produced are reported below.

A borosilicate glass sphere (Thermo fisher Scientific), with nominal radii R = 5 μm for nanoindentation (see **4.3.3**) and R = 10 μm for adhesion spectroscopy (see **4.3.1**), is attached to a tipless silicon cantilever (Micromash HQ:CSC38/tipless/no Al, spring constant *k* = 0.02 - 0.03 N/m) exploiting capillary adhesive forces or a small amount of vaseline as temporary adhesive. The glass sphere is then covalently attached to the silicon cantilever by thermal annealing for two hours in a high-temperature oven at the softening point temperature (T=780 °C). The resulting monolithic colloidal probe can be washed/cleaned aggressively after use in order to remove contaminants. The characterisation of the CP radius is performed by atomic force microscopy (AFM) reverse imaging of the probe on a spiked grating (TGT1, Tips Nano), as detailed in Indrieri *et al*.^93^. The spring constant is calibrated using the thermal noise method^95,96^ where special correction factors are applied in order to take into account the relevant dimension and mass of the glass sphere^94,97^.

### 4.2 Production of nanotopographical probes and substrates

For the production of the nanotopographical CPs (for adhesion force spectroscopy) and for the nanostructured substrates (for the retrograde actin flow experiments), ns-ZrO_2_ films are deposited on the colloidal probes, or glass substrates, respectively, exploiting the Supersonic Cluster Beam Deposition (SCBD) technique (described in detail in the Ref.s^26,98,99^). Briefly, SCBD allows the production of nanostructured surfaces characterised by a disordered topography, yet with accurately controllable and reproducible morphological properties such as rms roughness, cluster size distribution and porosity^100,101,102^. Partially oxidised zirconia clusters are produced within the cluster source and then extracted into the vacuum of the expansion chamber through a nozzle to form a seeded supersonic beam. Clusters are collected directly on the CPs, or glass substrates, intercepting the beam in the deposition chamber. Upon landing on the sample surfaces, clusters form a nanostructured, highly porous, high-specific area, biocompatible ns-ZrO_2_ film^27,32^. The protocol for the functionalisation of the colloidal probes with the nanotopographical surfaces achieved by zirconia cluster-assembling is detailed in Chighizola *et al*.^16^. The specific surface area, the rms roughness (defined as the standard deviation of the height of the surface, 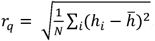, where *h*_*i*_ is the height of the surface in a specific point, N is the number of pixels of the map and 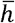 is the average height), the average lateral dimensions of the largest morphological features (the correlation length ξ) typically increase with ZrO_2_ film thickness^100,102,103^.

### 4.3 AFM experiments

All the AFM experiments have been performed using a Bioscope Catalyst AFM (Bruker), mounted on top of an inverted optical microscope (Olympus X71). Isolation of the system from noise was obtained by means of both, an active anti-vibration base (DVIA-T45, Daeil Systems), and an acoustic enclosure for the AFM (Schaefer, Italy). During the AFM measurements performed on live cells, the temperature of the cell culture medium was maintained at 37°C using a perfusion stage incubator (PSI, Bruker) and a temperature controller (Lakeshore 331, Ohio, USA). Before every measurement the deflection sensitivity was calibrated *in situ* and non-invasively by using the previously characterised spring constant as a reference, according to the SNAP procedure described in Schillers *et al*.^104^ and based on the assumption that the correct deflection sensitivity value must provide, through the noise calibration, the reference value of the spring constant. The cell regions to be investigated (portion of the membrane to be scanned or location of the force curves to be acquired) were chosen by means of the optical microscope; the accurate alignment of the optical and AFM microscopes was obtained using the Miro (Bruker) module integrated in the AFM software. Data processing of the sets of curves and scan images was carried out in Matlab (Mathworks) environment using custom routines.

#### 4.3.1 Morphological characterisation of surfaces (substrates and cells)

Morphological images of nanostructured zirconia thin films deposited on colloidal probes were acquired with a Bioscope Catalyst AFM (Bruker), in tapping mode. NCHV probes from Bruker, with resonance frequency about 300 kHz, force constant k= 40 N/m, and nominal tip radius 10 nm, were used for these experiments. The topographic maps have been collected with a sampling resolution of 1−2 nm/pixel using as scan rate of 1 Hz.

For the morphological surface characterisation of the PC12 cells, the cells were fixed with 2% paraformaldehyde/0.1% glutaraldehyde in PBS for 30 min after 1 day of culturing on poly-L-lysine (PLL)-coated (0.1% w/v PLL solution (Sigma) for 30 min at RT) glass-bottomed plates (Willco Wells). The fixed cells were characterised using a Bioscope Catalyst AFM from Bruker. The AFM images (with scan size of 3-5 μm) were acquired in PBS buffer in peak-force tapping mode. Because of the significant height variations within the samples (up to several micrometres), cantilevers with a tip height of 14 μm were used (VistaProbes CS-10). The scan rate was set to 0.6 Hz, the ramp frequency at 1 KHz, and the sampling resolution was set to 4096 × 1024 points. To avoid degradation of the glycocalyx, the measurements were performed within max. 24 hours following the fixation.

#### 4.3.2 Adhesion force spectroscopy and data analysis

Adhesion force spectroscopy measurements were performed as discussed in detail in Chighizola *et al*.^16^. The nanotopographical colloidal probes (R= 10 μm) were incubated with the cell culture medium (Ø phenol red) for >30 min at 37°C before the actual measurements (to guarantee absorption of ECM proteins, such as fibronectin and vitronectin, present in the serum to the probes). Sets of force-distance curves were acquired at locations on the cells’ body selected by means of the optical microscope. Force curves (FCs) containing 8192 points each were recorded on live PC12 cells, with ramp length *l* = 10 μm, maximum load *F*_*max*_ = 1 nN and retraction speed at *v*_*r*_ = 16 µm/s. The pulling speed was kept relatively low to reduce hydrodynamics effects^105^. To measure the early steps of cellular adhesion in the critical phase of nascent adhesion formation towards FA maturation, we selected five contact times (*cts*): 0, 20, 60, 120, 240 s, accordingly. During the contact time, the Z-piezo position was kept constant using the Z closed-loop feedback mode. The data analysis of the retraction curves was carried out using the protocol presented in Chighizola *et al*.^16^.

#### 4.3.3 Glycocalyx characterisation by AFM nanoindentation

##### Glycocalyx thickness

Nanomechanical indentation experiments were performed using colloidal probes with radius R = 5 μm following the approach presented by Puricelli *et al*.^106^. To get quantitative information about the thickness of the glycocalyx, we applied the brush model implemented by Sokolov *et al*.^34,35^. This model was developed with the purpose to decouple the single mechanical contributions of the glycocalyx compression and the underlying cell deformation, allowing isolating the force needed to compress the glycocalyx (for details, see **SI2** and **SI – Fig. S2-3**).

##### Glycocalyx stiffness

Mechanical properties of the glycocalyx were calculated following the approach described by Wiesinger *et al*.^107^. The effective glycocalyx spring constant or stiffness, defined following the Hook’s Law *K*_*GC*_ = ^*dF*^/_*dh*_ has been extracted from each FC by selecting a suitable small region where the entropic resistance to compression^34^ can be considered linear.

#### 4.3.4 Statistics and error analyses

For each observable *ψ*_*FC*_ extracted by each force curve (FC) (*e*.*g*., the adhesion force F_a_, the work W, the GC stiffness K_GC_), a mean value 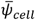 was evaluated for each cell.

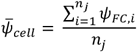

where nj is the number of FCs per each cell, j = 1,…N, *N* being the number of cells investigated for each condition.

The mean value 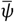 representative of the cell population in each condition was evaluated as:

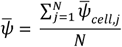

The effective error 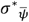 associated to 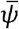 was obtained by summing in quadrature the standard deviation of the mean 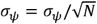 being the standard deviation of 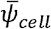values, and an estimated instrumental error 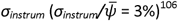^106^:

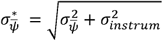

#### 4.3.5 Modelling of the surface accessibility in dependency of the cell membrane compliancy

The strategy used to determine the potential interacting interface of the cells in contact with nanotopographical surfaces, consists in analysing the morphological AFM maps (n = 3 per roughness value) of the cluster-assembled thin zirconia films (**Fig. 3A**), by applying masks at different interfacial heights (**Fig. 3B,C**).

The starting threshold in z was chosen from the value of the highest asperity of the morphological map, and we have decreased the subsequent thresholds by 5 nm per step (as schematically shown in Figure 3A). For each threshold used, we mask the portion of the AFM image below the chosen threshold. For each threshold value, we calculated the number of asperities, the mean surface area of asperities, the sum of the 3D surface area of the asperities identified on the surface (total surface area), the specific surface area (defined as the ratio of 3D area of the asperities and their projected 2D area), and the coverage (defined as the sum of the 2D projected area of the asperities identified over the entire 2D area).

In order to integrate the mechanotransductively-relevant parameter integrin ligand spacing, we have calculated the change in the number of zirconia asperities at each threshold by expanding the perimeter of the 2D area of each asperity of 30 nm in x-y direction, since 60 nm was identified as critical ligand spacing distance for the maturation of integrin adhesion complex^42,43,44^. Converging asperities were considered as connected adhesion areas and counted as a single entity. Furthermore, it was determined that, at least, four integrin binding sites in a vicinity of 60 nm can serve as a minimal adhesion unit which promotes IAC maturation^46,47,48,45^. For this reason, we analyzed also the number of asperities at different interfacial depths, characterised by a 3D surface area of, at least, 60 nm x 60 nm = 3600 nm^2^.

### 4.4 Cell culture and preparations for experiments

The experiments were performed with the neuron-like PC12 cells (PC12-Adh clone, ATCC catalogue no. CRL-1721.1TM). This cell line has shown to respond to the nanotopographical cues provided by the substrates produced by SCBD (in particular ns-Zr15) by mechanotransductive processes and signalling, which we have profoundly characterised in previous studies^9,10,16^.

For routine cell culture (subculturing every 2-3 days using trypsin/EDTA solution), the PC12 cells were maintained (in a humid atmosphere at 37°C and 5% CO_2_) in an incubator (Galaxy S, RS Biotech) provided with RPMI-1640 medium (supplemented with 10% horse serum, 5% fetal bovine serum, 2 mM 1-glutamine, 10 mM HEPES, 100 units mL^-1^ penicillin, 100 µg mL^-1^ streptomycin, 1 mM pyruvic acid). All reagents and materials from Sigma, also in the following, if not stated otherwise.

For the experiments, the cell detachment was done on the day before with an EDTA (1mM in PBS) solution to not digest the protein component of the glycocalyx. The cells were counted with an improved Neubauer chamber before plating them in a low concentration (4000 cells per cm^2^, to have mostly single, separated cells) on the indicated substrates. The cells were then maintained in the incubator overnight to guarantee good cell adhesion.

For the experiments involving AFM measurements, the cells were seeded the day before the experiment on Ø 40 mm glass-bottomed plates (Willco Wells) that were coated before with PLL (0.1% w/v PLL solution for 30 min at RT). Solutions without phenol red were used, as it results harmful for the AFM tip holder.

For the actin dynamics measurements, one day before the experiment the cells (after LifeAct™ transfection, see 4.6.1) were instead plated on Ø 35 mm glass-bottomed dishes (MatTek, the Ø 13 mm glass was in the centre of the plate) on which either flat zirconia film (produced by ion gun sputtering) or nanostructured cluster-assembled zirconia film (fabricated by SCBD) have been deposited.

In case of glycocalyx-targeting enzymatic treatment, an enzyme cocktail was applied to the cell culture medium to obtain a final concentration of 13.5 U/mL hyaluronidase II, 150 mU/mL chondroitinase ABC, 300 mU/mL heparinise III and 150 mU/mL neuraminidase (these concentrations were inspired by the work of Zeng et al.^108^). The cells were incubated with these enzymes for 2 hours right before the experiments. Following the incubation, the cell culture medium was discarded. After washing 3x with PBS (to remove the digested glycocalyx), fresh medium was added containing again the enzymatic cocktail to avoid reassembly of the glycocalyx during the measurements.

For the Y27632 inhibition against ROCK, the cells were pre-incubated with the inhibitor 15 min before the measurements. The inhibitor was used at a final concentration of 10 µM.

### 4.5 Glycocalyx characterisation by wheat germ agglutinin staining and 3D-Structural Illumination Microscopy (3D-SIM)

The glycocalyx of live PC12 cells in the indicated experimental conditions (prepared as described in 4.4) was marked 30 min before the recording by adding Oregon Green™ 488 conjugate wheat germ agglutinin (WGA, Thermofisher) to the medium at a final concentration of 25 µM/mL. WGA binds sialic acid and N-acetylglucosaminyl residues. After the 30 min incubation (in the incubator at 37°C and 5% CO_2_), the cells were washed 3x with PBS to remove the excess of WGA (to reduce the background fluorescence noise) and new phenol red-free medium (prewarmed to 37°C) was given to the cells (in case of glycocalyx-targeting enzymatic treatment containing again the enzymes). Then the plates were immediately transferred to the 3D-SIM Nikon A1 microscope (equipped with a stage-top incubation system (OkoLab), conditions: humid, 37°C, 5% CO_2_) to take the images. An oil immersion 63x objective was used.

The thickness of the glycocalyx was determined from crops of the cell borders in the plane just above the cell/substrate interaction zone processed by means of ImageJ/FIJI^109^ (for better distinction the “Enhance Contrast” tool was applied). The thickness measurements from the cropped images were then done with a customised macro in the MatLab environment that automatised the data collection after marking the thickness with lines. The graph was obtained with the help of Plotsofdata web apps^110^.

### 4.6 Visualisation, recording and quantification of the actin dynamics

#### 4.6.1 Transfection with LifeAct^™^ and recording

To visualise the actin dynamics, the PC12 cells were transfected with mCherry-LifeAct (the vector was a kind gift by Prof. Giorgio Scita). For this purpose, the cells were plated on Ø 35 mm petri dishes (TRP) at a low confluency (10-15%) the day before the transfection. At the beginning of the next day, the transfection was done with Lipofectamine LTX® performing the following steps:

- Solution 1: 6 µl of Lipofectamine LTX® was added to 150 µl RPMI base medium (incubation for 5 min at RT).
- Solution 2: 2 µl of the mCherry-LifeAct vector DNA (concentration 1 µg/µL) was added to 150 µl RPMI base medium, then 2 µl Plus™ reagent was added.
- The two solutions were mixed and incubated for 20 min at RT and then added dropwise to the cells in a homogeneous way.

The success of the transfection was controlled the next day with a Zeiss Axiovert 40 CFL with an epifluorescence module.

After transferring the transfected cells onto the indicated substrates (see 4.4), the actin dynamics (before and after glycocalyx removal, see 4.4) were then recorded the next day with a Nikon A1R confocal microscope at the UNI^TECH^ NOLIMITS Imaging facility of the University of Milan. The microscope is equipped with a stage-top incubation system (OkoLab) to guarantee humidification, 37°C and 5% CO_2_ during the live cell imaging. An oil immersion 60x objective was used. The recordings were done with a frame rate of 0.5 frames/sec for 2 minutes.

#### 4.6.2 Analysis of the actin dynamics by Particle Image Velocimetry

Particle image velocimetry (PIV) was performed to quantify the actin dynamics. This analysis results in a vector field, highlighting the local movement of the actin fibres in time, from which the average flow speed can be obtained.

##### Pre-processing

Photobleaching was corrected in Fiji hypothesising linear decay of the signal and the cell area was segmented by thresholding. A temporal resolution of 6 s between frames was used (intermediate frames were removed).

##### Analysis

PIV analysis is described in detail elsewhere^111^. Briefly, a region of interest (source box) is searched for in a larger area (search box) of the subsequent frame by two-dimensional cross-correlation. A grid of regions of interest is drawn over the image, to obtain a full local tracking of the actin filaments. For this analysis, the PIV parameters were optimised as follows: source box 0.6 µm, search box 1 µm, distance between regions of interests (grid) 0.4 µm, correlation coefficient threshold of 0.3. Spatial and temporal convolutions are subsequently performed to interpolate the vector field. The defined kernels were 5 µm (sigma = 1 µm) and 30 s (sigma = 12 s), respectively. Colour maps display flow velocity within a 0-2 µm/min scale.

## Supporting information

PIV video flat-Zr

PIV video flat-Zr plus enzymatic cocktail

PIV video ns-Zr15

PIV video ns-Zr15 plus enzymatic cocktail

## ACKNOWLEDGEMENTS

We acknowledge the support of the European Union’s Horizon 2020 research and innovation programme under the Marie Skłodowska-Curie grant agreement No. 812772, project Phys2Biomed, and under FET Open grant agreement No. 801126, project EDIT. BS and SM are funded by the European Research Council (ERC) under the European Union’s Horizon 2020 research and innovation programme (grant agreement No. 681808). PM and CS acknowledge support from the European Union FP7-NMP-2013-LARGE-7 “Future NanoNeeds” programme. CL gratefully acknowledges funding from Miur – PRIN2017, Prot. 2017YH9MRK. We thank the UNI^TECH^ NOLIMITS Imaging facility of the University of Milan (in particular Dr. Miriam Ascagni) for support in regard to the realisation of the experiments involving confocal microscopy and 3D-SIM. We thank Prof. Giorgio Scita, Prof. Jan de Boer, Prof. Johanna Ivaska, Dr. Martina Lerche and Aleksi Isomursu for critical reading of the drafts and valuable comments.

## AUTHOR CONTRIBUTIONS

Conceptualisation: MC, AP, CS; methodology – probe fabrication and characterisation: MC, APr, FB, CP; methodology – SCBD: APr, CP; methodology – modelling: MC, FB, CS, AP; methodology – cell culture and preparation: TD, MDU, CS; methodology – glycocalyx characterisation AFM: MC, LC, TD, CS, AP; methodology – AFM adhesion force spectroscopy: MC, TD, AP; methodology – actin dynamics: MDU, CS; methodology – glycocalyx characterisation 3D-SIM: MDU, CS; data curation and formal analysis – modelling: MC, FB, AP; data curation and formal analysis – AFM glycocalyx characterisation: MC, LC, TD, CS, AP; data curation and formal analysis – AFM adhesion force spectroscopy: MC, TD, CS, AP; data curation and formal analysis – actin dynamics: MDU, SM, BS, CS; data curation and formal analysis - glycocalyx characterisation 3D-SIM: CF, CS; original draft writing: MC, CS; draft reviewing and editing: MC, TD, FB, CL, PM, SM, BS, AP, CS; supervision: CL, PM, BS, CS, AP; resources, funding and project administration: CL, PM, BS, AP. Author contributions were allocated adopting the terminology of CRediT – Contributor Roles Taxonomy.

