## Supplementary Information for "The glycocalyx affects the mechanotransductive perception of the topographical microenvironment"

**of topographical microenvironment**

**^1^Matteo Chighizola^F^, ^1,2^Tania Dini, ^3^Stefania Marcotti, ^1,4^Mirko D’Urso, ^1^Claudio Piazzoni, ^1^Francesca Borghi, ^1^Anita Previdi, ^1^Laura Ceriani, ^1^Claudia Folliero, ^3^Brian Stramer, ^1^Cristina Lenardi, ^1^Paolo Milani,**

**^1^Alessandro Podestà*, ^1^Carsten Schulte***

**Affiliations:**

^1^Interdisciplinary Centre for Nanostructured Materials and Interfaces (C.I.Ma.I.Na.) and Department of Physics “Aldo Pontremoli”, University of Milan, Milan, Italy

^2^The FIRC Institute of Molecular Oncology (IFOM), Milan, Italy

^3^Randall Centre for Cell and Molecular Biophysics, King’s College London, London, United Kingdom

^4^Department of Biomedical Engineering, Institute for Complex Molecular Systems, Eindhoven University of Technology, Eindhoven, Netherlands

^F^ first author

***Co-last and corresponding authors:**

Carsten Schulte:

Alessandro Podestà:

**Keywords:**

Mechanotransduction, glycocalyx, integrin adhesion complexes, molecular clutch, force loading, focal adhesion, nanostructured cell microenvironment, nanotopography, Atomic Force Microscopy, colloidal probes.

### 1. Implementation of the Sokolov’s brush-Model

Following the approach described in Refs ^1,2,3^, the cell and the glycocalyx can be thought of as a two-layer system with two different elastic properties. In the geometry of the indentation experiment the ***probe-membrane*** distance *h* can be described as:

$h=Z-Z_{0}+i+d$	(1)

Where *Z* is the relative piezo position, *Z_0_* is the undeformed position of the cellular membrane, *d* the deflection of the cantilever, which is positive when the cantilever is bent upward (in repulsive regime), and *i* the indentation calculated using the Hertz model

$i=\left. \left( \frac{3k(1-v^{2})}{4E}R^{\frac{1}{2}} \right) \right.^{\frac{2}{3}}d^{\frac{2}{3}}$	(2)

Here *E* is the Young’s modulus of the cell body, *k* is the spring constant of the probe, *v=0.5* is the Poisson coefficient.

Assuming that the glycocalyx layer is much softer than the cell body, the cantilever reaches an applied force where the glycocalyx can be considered completely squeezed (*h=0*) before reaching the maximum load set. This assumption depends obviously on the value of the force threshold applied, which should be sufficiently large; in this condition $\left( Z-Z_{0} \right)+d=-i$, as in conventional indentation experiments.

The force needed to completely squeeze the glycocalyx cannot be known *a priori* and must be found. Our approach was to divide the total effective deformation δ*_eff_* (squeezing of the glycocalyx plus cellular deformation) in intervals *s=200* nm and then apply the hertz model in those intervals. The idea behind this approach (already used to detect the glycocalyx mechanical response^4,5^) is that the first mechanical response perceived by the probe would be the resistance to compression of the pericellular brush, then applying more and more force the glycocalyx will squeeze and progressively transfers rigidly the applied force to the cell below. Once the glycocalyx will be completely compressed the resulting effective Young’s modulus *E_eff_* will stabilise to the cell body-bulk values. The *E_eff_* was plotted as function of the δ*_eff_* and then we looked at discontinuity in the *E_eff_* and where it reaches a plateau. We interpreted the plateau as a situation where the glycocalyx is fully compressed and what is measured is the Young’s modulus of the cell body. The result of this analysis was made on a population of ~20 cells in standard conditions (represented in **SI - Fig. S1**).

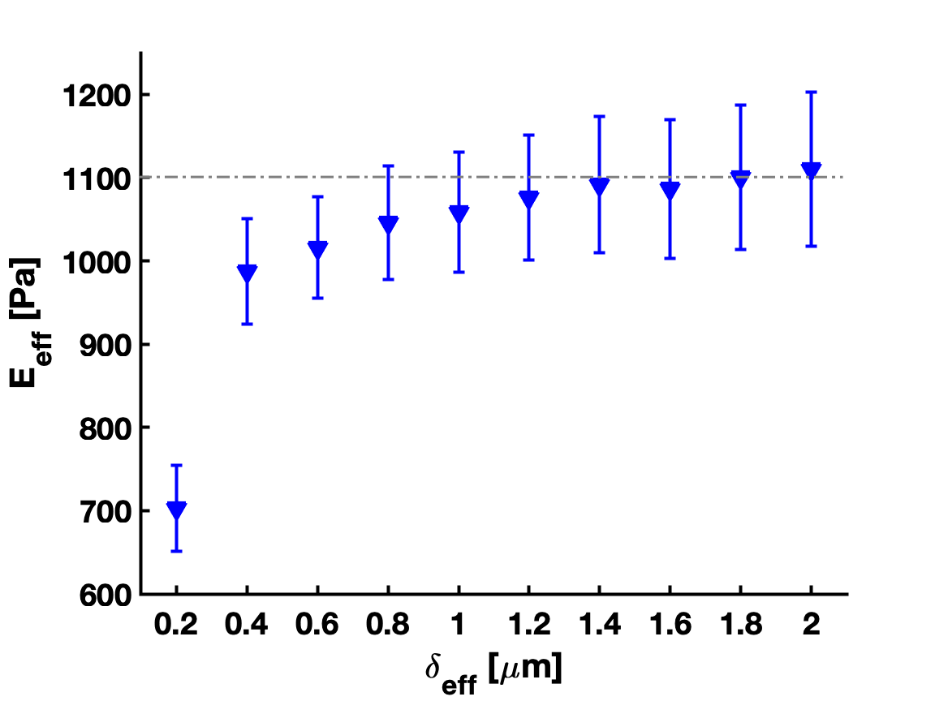

***SI – Fig. S1.*** *Effective YM E_eff_ evaluated per each intervals s as a function of the effective indentation d_eff._ Data represent the mean YM averaged over the cell population, while the error bars represent the standard deviation.*

It can be noted that after the first 200 nm a sudden increase of the *E_eff_* is present. The subsequent smooth increase of *E_eff_* can be attributed to inhomogeneity in the elastic properties of the cellular interior (*e.g.*, cortex vs nucleus) and to finite-thickness effects^6^. The corresponding force threshold, at which the glycocalyx is fully compressed was set at *F_thr_= 0.1-0-2 nN.*

Fitting the hertz model in a region just above *F_thr_* (where $h\cong0$) would provide a value of *E* in the superficial region. The intersection of a back extrapolation of the hertz model and the horizontal line (*F=0*) would give the value of the parameter *Z_0_*, that represents the contact point between the probe and the cell membrane*.* Using then Eq. 2, it is possible to evaluate the indentation *i* (which actually is the real deformation of the cell, beneath the glycocalyx compression). With all these parameters, it is possible to calculate the probe-membrane distance *h* at every position of the piezo.

Note that care must be taken in the choice of the indentation range for the fit of the Hertz model; as it is evident from **SI - Fig. S1** that the inner cell body is anything but uniform. The cell body is isotropic and this inhomogeneity is reflected by the change of the YM. When choosing a fitting region, one has to take into account that, looking at Eq. 1, the aim of the procedure is to quantify the cellular deformation within the effective indentation δ*_eff_*, in order to obtain the residual deflection of the probe, *i.e.* the effect only due to the brush steric resistance. Therefore, the YM should be extracted from the first indentation range above the threshold, identified in **SI - Fig. S1**, otherwise the probe-membrane distance and then the glycocalyx thickness would be estimated inaccurately.

Once the probe-membrane distance is evaluated the FCs must be rescaled on this new distance as shown in **SI - Fig. S2**.

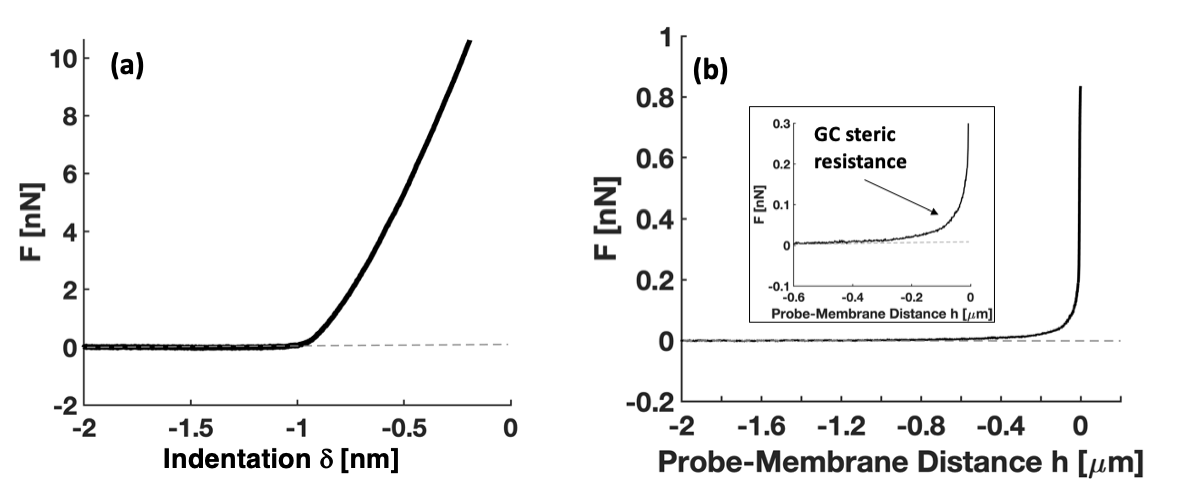

***SI – Fig. S2. (a)*** *Original Force Vs Indentation curve taken over the cell body.* ***(b)*** *Same curve rescaled over the new distance h. After the glycocalyx complete compression (F = 0.2 nN), the probe is in contact with the membrane and the rest of the curve collapses on the h=0 vertical axis. The inset shows the appearance of the glycocalyx’ steric resistance a few hundred of nm before the probe gets into contact with the cell membrane.*

Because of the specific force dependence and its physical nature (actually intertwined polymer chains) the glycocalyx layer cannot be described by an effective YM. The glycocalyx layer increases its stiffness during the compression. To describe the glycocalyx behaviour quantitatively, the Alexander-de Gennes model was used, implemented by Butt *et al.*^7^. It describes the steric interaction between a spherical indenter of radius *R* and a flat surface due to the existence of superficial *entropic brush* ^8^:

$F\left( h \right)=50k_{B}TRN^{\frac{3}{2}}{L e}^{\left( -2\pi\frac{h}{L} \right)}$	(3)

Where *k_b_* is Boltzmann’s constant, *T* is the medium temperature, *R* is the probe radius, *N* the brush grafting density and *L* the brush length-thickness. From the fitting of the force rescaled data with Eq (3), it is possible to extract the thickness parameters *L.*

### 2. 3D-SIM Glycocalyx imaging

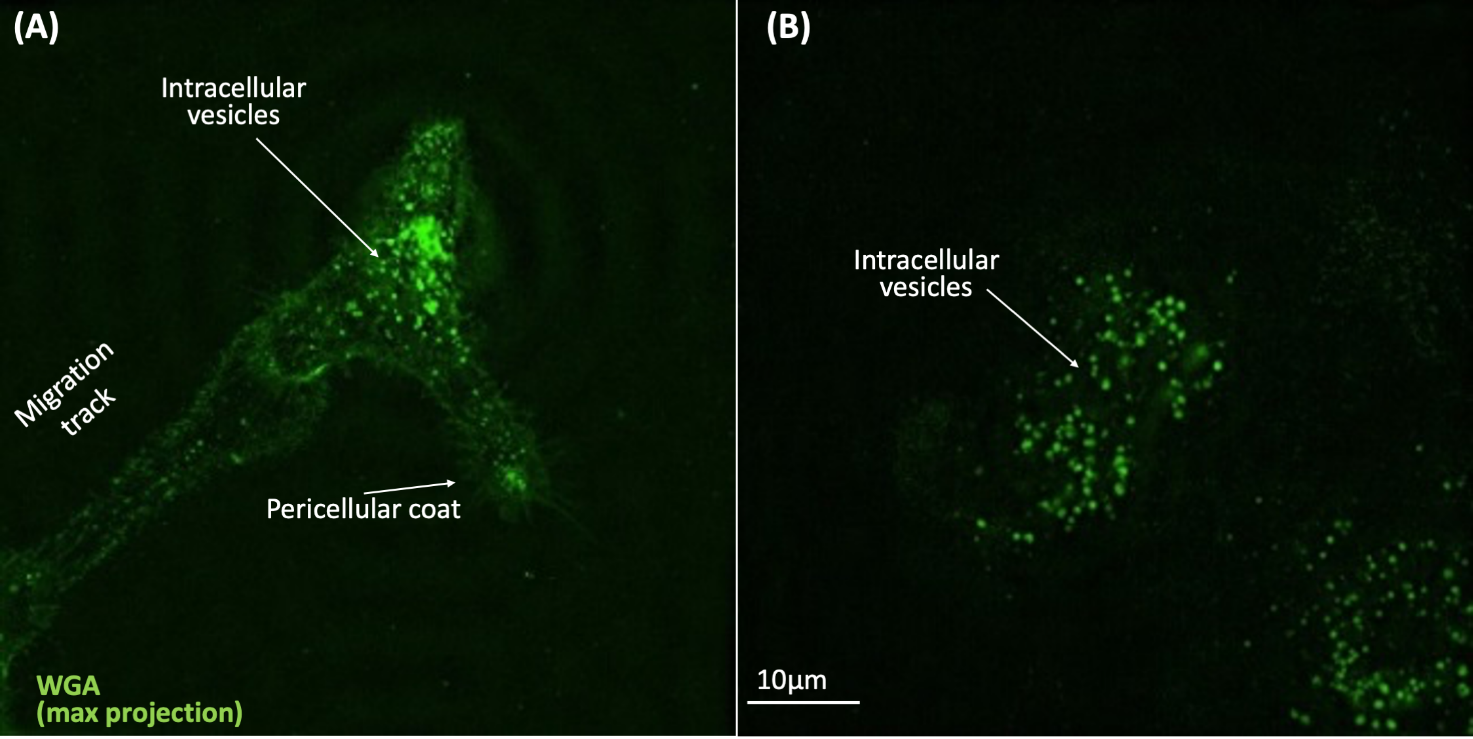

***SI – Fig. S3.*** *Representative 3D-SIM images (max. projection) of* ***(A)*** *PC12 on PLL, native or* ***(B)*** *treated with the glycocalyx-targeting cocktail, marked with Oregon Green^®^488 conjugated wheat germ agglutinin (WGA) which binds to sialic acid and N-acetylglucosaminyl residues.*

The native PC12 cells show (**Fig. SI-S3A**) different glycocalyx containing (extra)cellular structures stained in green, *i.e.* intracellular vesicles filled with glycocalyx components, the pericellular glycocalyx coat and a migration track/migrasome (an extracellular structure/organelle known to be left behind some migrating cells^9,10,11^). After the enzymatic treatment (**Fig. SI-S3B**) all staining disappeared, except for the intracellular vesicles.

### 3. Representation and analysis of the nanotopographical substrate features

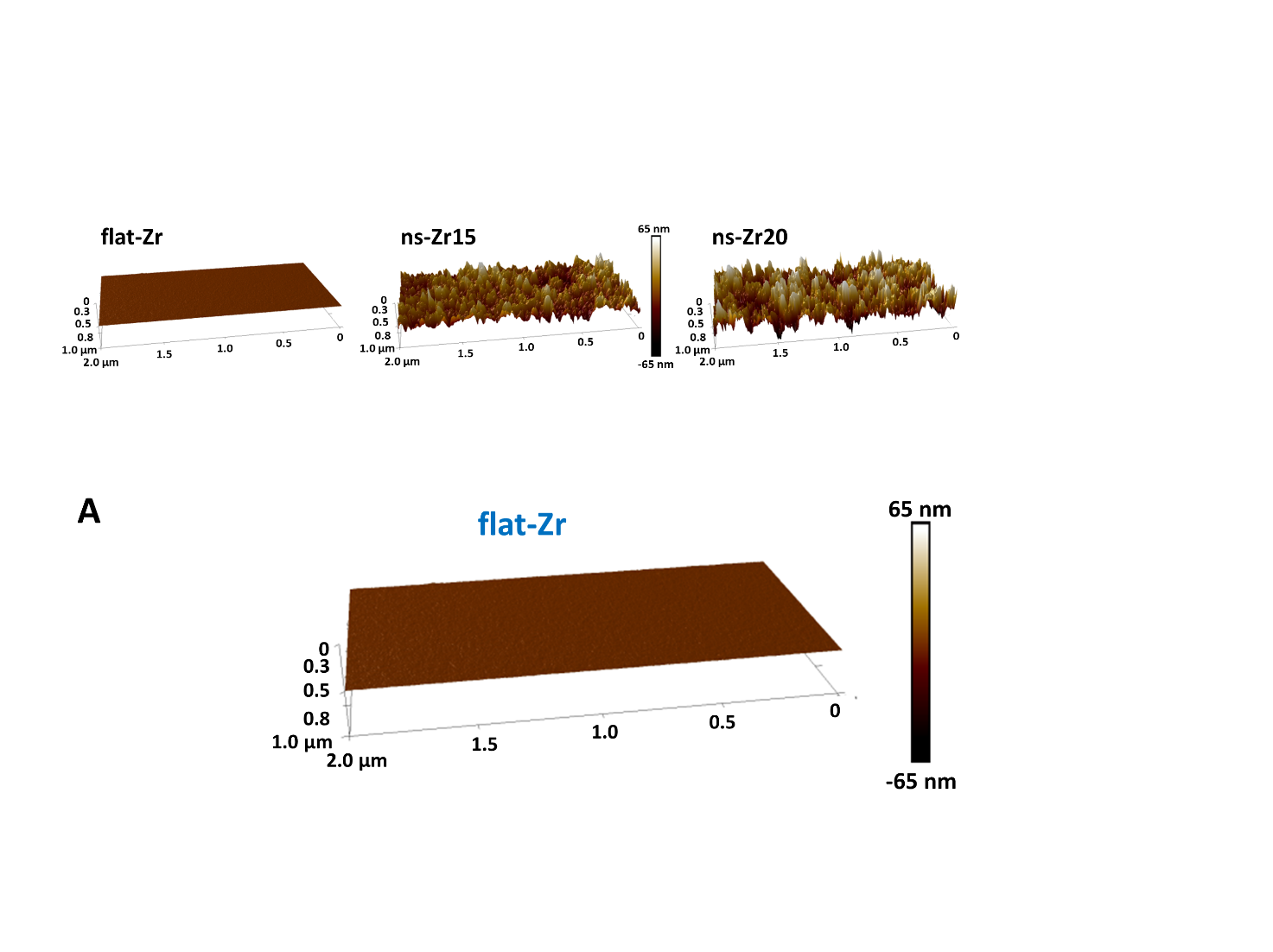

***SI – Fig. S4)*** *A representation of flat-Zr produced by Ion Gun sputtering.*

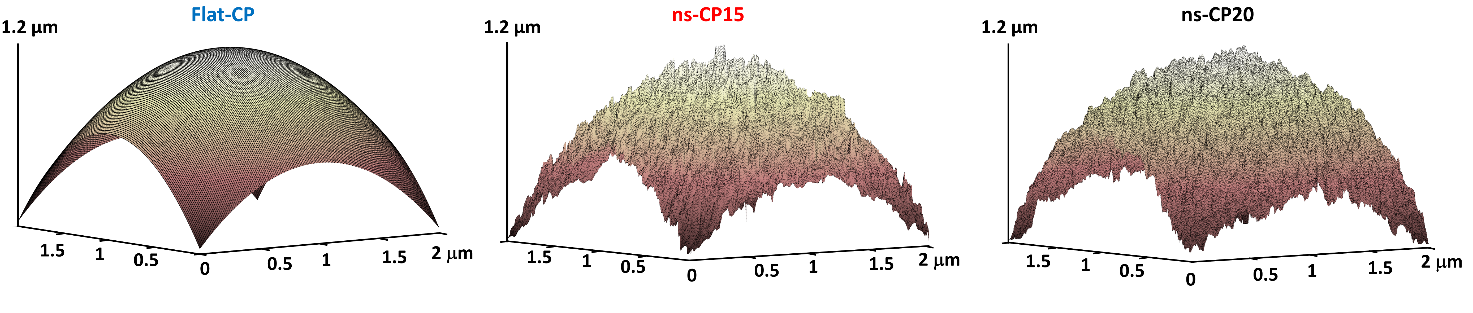

***SI – Fig. S5)*** *AFM scans of colloidal probes (CP) decorated with zirconium oxide (ZrO_x_). Left, CP with flat thin film of ZrO_x_ deposited with an ion gun. Middle and right, CPs decorated with nanostructured ZrO_x_ deposited by supersonic cluster beam deposition. Middle, CP with R_q_ = 15 nm root-mean-square (rms). Right, CP with R_q_ = 20 nm rms. The images were taken over probes with 5 μm radius to appreciate both, the nanostructure and the sphere curvature.*

### 4. Further parameters extracted from the adhesion force spectroscopy measurements – Work W and Number of tethers N_t_

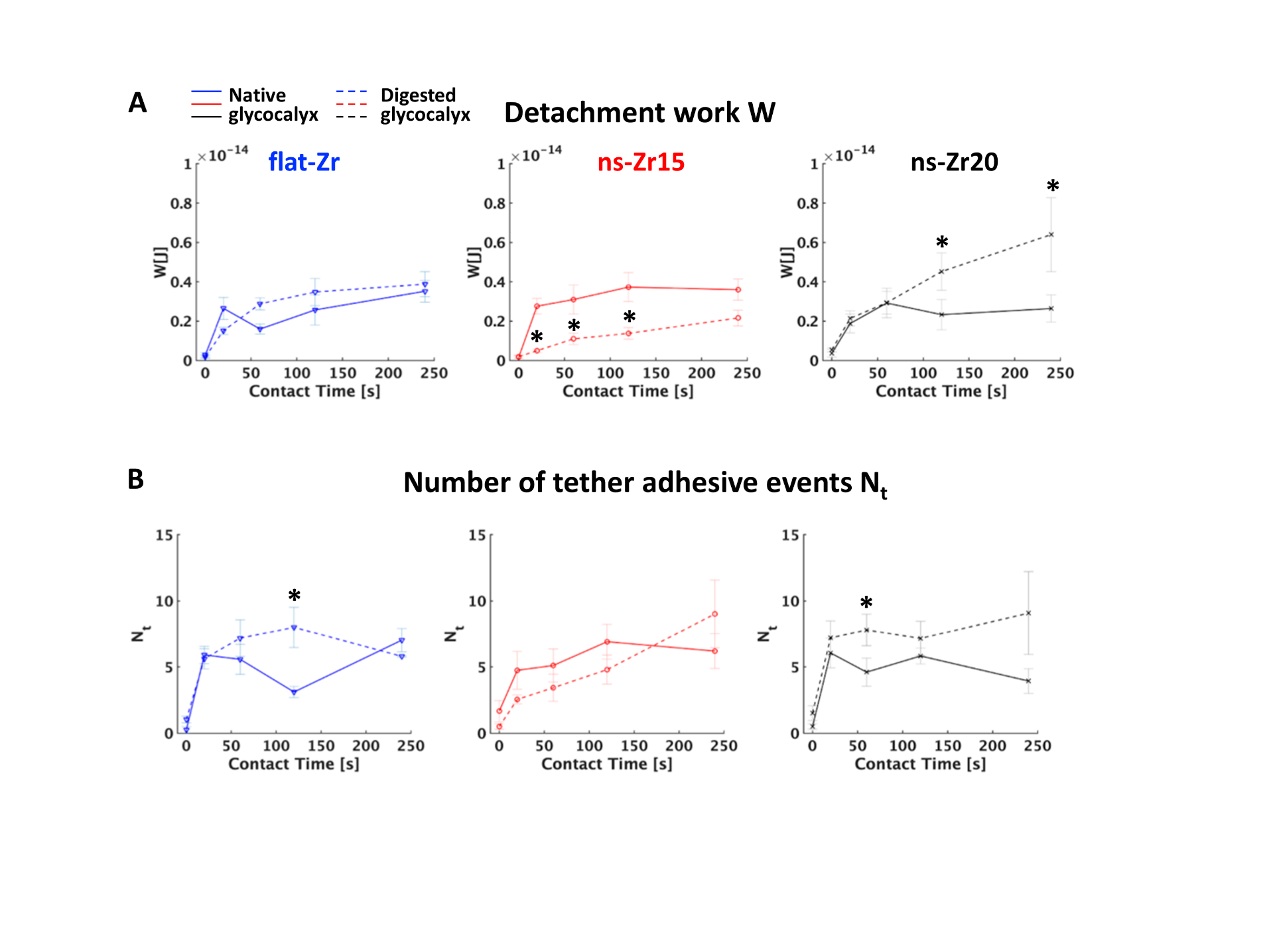

***SI – Fig. S6)*** *The panel shows the results of the adhesion force spectroscopy measurements for the different indicated experimental condition for the parameters* ***(A)*** *Detachment work W and* ***(B)*** *Number of tether adhesive events. Asterisks indicate a statistically significant difference between untreated and enzymatic treatment condition.*

### 5. Jump force data distribution

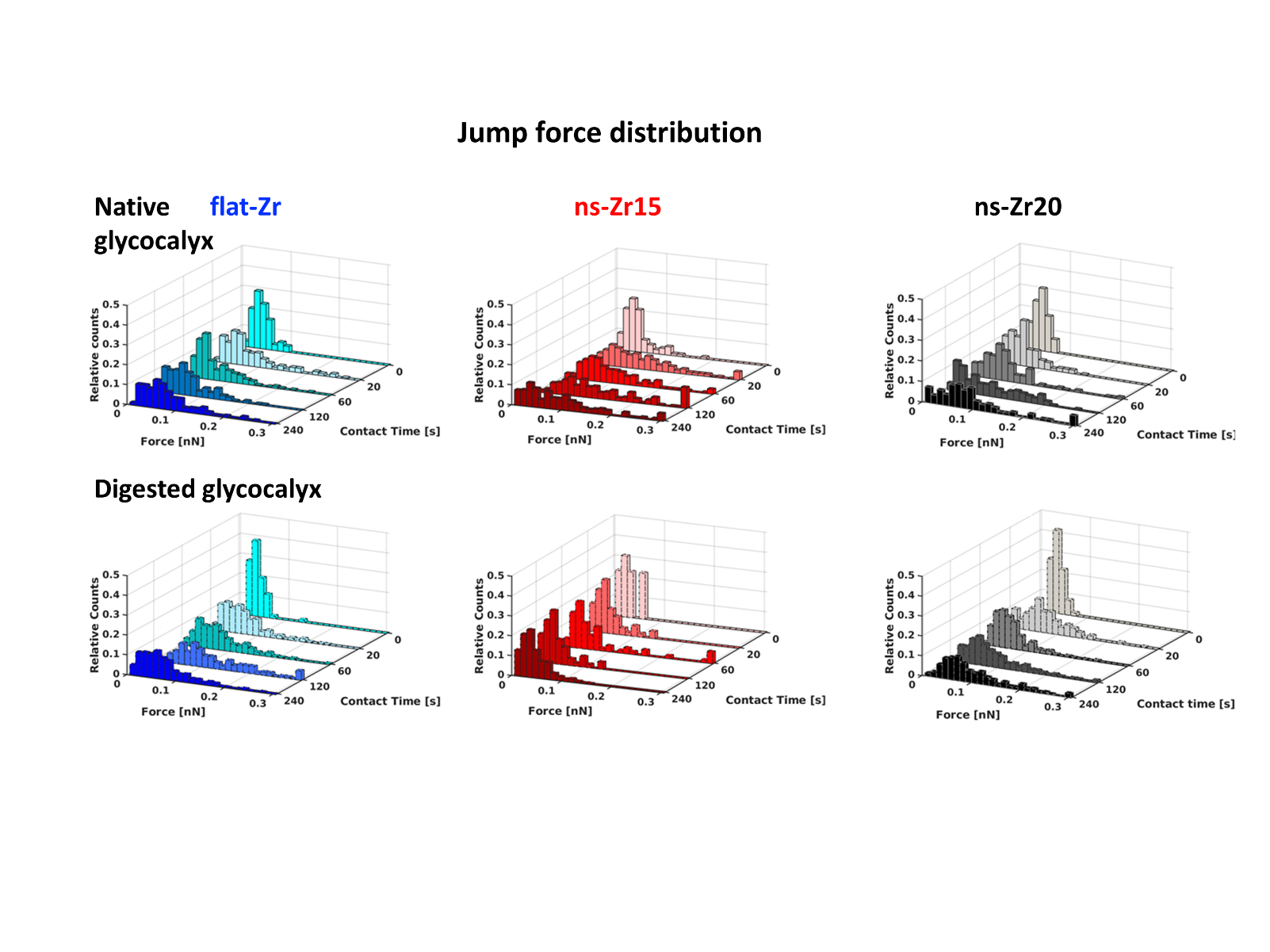

***SI – Fig. S7)*** *The panel shows the jump force distribution results in the indicated different experimental conditions of* ***Fig. 3D*** *in a finer binning to highlight the stronger dispersion of the higher forces in the untreated PC12 on ns-Zr15.*
